# Correlating phylogenetic and functional diversity of the *nod*-free but nodulating *Bradyrhizobium* phylogroup

**DOI:** 10.1101/2023.06.14.544914

**Authors:** Lu Ling, Alicia Camuel, Sishuo Wang, Xiaojun Wang, Tianhua Liao, Jinjin Tao, Xingqin Lin, Nico Nouwen, Eric Giraud, Haiwei Luo

**Affiliations:** Simon F. S. Li Marine Science Laboratory, School of Life Sciences and State Key Laboratory of Agrobiotechnology, The Chinese University of Hong Kong, Shatin, Hong Kong SAR, China; Shenzhen Research Institute, The Chinese University of Hong Kong, Shenzhen, China; IRD, Laboratoire des Symbioses Tropicales et Méditerranéennes (LSTM), UMR IRD/Institut Agro/INRAE/ Université Montpellier France; PHIM Plant Health Institute, Université Montpellier, IRD, CIRAD, INRAE, Institut Agro, Montpellier, France; Institute of Environment, Energy and Sustainability, The Chinese University of Hong Kong, Shatin, Hong Kong SAR, China

**Keywords:** *Bradyrhizobium*, *Aeschynomene*, nodulation, population genomics

## Abstract

*Bradyrhizobium* is a main rhizobial lineage of which most members nodulate legume plants using Nod factors (NFs) synthetized by the *nod* genes. However, members of the Photosynthetic supergroup (phylogroup) within *Bradyrhizobium* (PB) are *nod*-free but still capable of establishing nitrogen-fixing nodules with some tropical legumes of the *Aeschynomene* genus.

These unusual findings are based on the genomic sequences of only 13 PB strains, and almost all were isolated from *Aeschynomene* nodules. Here, we investigate the diversity of *Bradyrhizobium* in grassland, forest, and rice field by *rpoB* amplicon sequencing and report that PB is mainly associated with rice root and rhizosphere. Moreover, we sequenced 209 new PB members isolated mostly from the rice field. The extended PB supergroup comprises three major clades: a basal clade with significant expansion of its diversity, followed by an intermediate clade composed by two strains, and a new clade exclusively represented by our new strains. Although the PB strains universally lack the canonical *nod* genes, all 28 assayed strains covering the broad diversity of these clades induced nodules on *Aeschynomene indica*. Interestingly, the three clades displayed significant differences in the efficiency of symbiosis, aligning well with their phylogenetic branching order. Our strain collection expands the ecological, phylogenetic and functional diversity of *nod*-free but nodulating *Bradyrhizobium*. With this expanded diversity, we conclude that the NF-independent nodulation of *Aeschynomene* is a common trait of this supergroup, in contrast to the photosynthetic trait originally thought as its unifying feature.

## Introduction

The genus *Bradyrhizobium* is one of the largest and most diverse rhizobial genera and the primary symbiont of a wide range of legumes (Parker, 2015; Sprent et al., 2017). It encompasses seven phylogenetic supergroups (phylogroups) (Avontuur et al., 2019; Ormeno-Orrillo and Martinez-Romero, 2019), among which the Photosynthetic *Bradyrhizobium* (PB) supergroup is one of the most special as photosynthesis is a rare trait in rhizobia (Avontuur et al., 2019).

Members belonging to this supergroup engage in a mutualistic relationship with some tropical semi-aquatic species of the *Aeschynomene* genus (Giraud and Fleischman, 2004), and they form nodules not only on roots but also on stems (Chaintreuil et al., 2013; Mornico et al., 2011).

Further, none of the completely sequenced PB members carry *nodABC* genes encoding the enzymes synthesizing the core structure of the nodulation factors (NF), the only exception being *Bradyrhizobium* sp. ORS285, which can use both a *nod*-dependent and -independent pathway reliant to the *Aeschynomene* host plant (Bonaldi et al., 2011; Giraud et al., 2007). This indicates that the PB phylogroup uses a novel mechanism of interaction with legume hosts that differs from the traditional universal NF-based pathway (Lerouge et al., 1990; Oldroyd, 2013). Moreover, all known PB strains carry photosynthetic genes except for four deep-branching members isolated from French Guiana as represented by *Bradyrhizobium* sp. STM3843 (Miché et al., 2010; Mornico et al., 2011).

Despite these unique features, the research on PB has been very limited. Currently, there are only 13 PB strains that have their genomes sequenced, largely limiting genome-based analysis on this important *Bradyrhizobium* phylogroup. Although most PB strains were isolated from *Aeschynomene* spp. nodules, the PB phylogroup was also detected in paddy soil, rice roots as well as in lake water (Chaintreuil et al., 2000; Okubo et al., 2013; Piromyou et al., 2015; Van Berkum and Eardly, 2002), suggesting that its members have much wider ecological niches than previously thought. Here, we report that rice fields are an important reservoir of PB among the terrestrial ecosystems through *rpoB* amplicon sequencing analysis. We isolated and genome-sequenced 209 isolates predominantly from three niches of rice (within root, rhizosphere, bulk soil) and discovered a novel deeply branching clade. We further show that phylogenomic diversity of PB members match well with their symbiosis efficiency with *Aeschynomene indica*.

## Materials and methods

All the methodological details were described in the Supplementary Text. Three plant species (*Oryza sativa* indica, *Houttuynia cordata,* and *Camphora officinarum*) and soils, each with three replicates, were collected from paddy field, grassland and forest, with each replicate 5-10 m apart from the others, respectively, in Hunan province, China (27.948 °N, 113.221 °E) in July 2021. Three typical niches (bulk soil, rhizosphere and root) of each plant species (three replicates) were each used for bacterial isolation using a modified arabinose-gluconate (MAG) medium. Basic soil characteristics were also measured. The taxonomic affiliation of isolates was determined by 16S rRNA gene analysis. Additionally, *Oryza sativa* japonica plants were collected in Hong Kong, China (22.418 °N, 114.080 °E) in May 2022 for *Bradyrhizobium* isolation following the same procedure.

DNA was extracted from fresh soil and plant root samples (0.25 g) using the DNeasy PowerSoil Pro Kit (QIAGEN) according to the manufacturer’s protocol and then sent to the company (Magigene, Guangdong) for *rpoB* amplicon sequencing. As the quality and quantity of DNA extracted from the second replicate of the rice root sample were poor and the amount of the root sample was insufficient to support multiple DNA extractions, amplicon sequencing was not performed on this sample. The quality control of the raw reads was performed with Trimmomatic v0.39 (Bolger et al., 2014). Subsequently, the paired-end reads of the *rpoB* amplicon sequences were processed with a denoising algorithm (DADA2) (Callahan et al., 2016) implemented in QIIME2 (Bolyen et al., 2019) to perform sequence denoising, dereplication, and chimera filtering to generate amplicon sequence variants (ASVs). The generated ASVs were filtered out the non-*rpoB* sequences and assigned to each *Bradyrhizobium* supergroup as well as each of the three clades of Photosynthetic supergroup through a phylogenetic placement method (Czech et al., 2022), which was used to determine their relative abundance.

To distinguish each *Bradyrhizobium* supergroup, two *rpoB* gene trees, using the full length (Fig. S1A) and amplified region (Fig. S1B) respectively, were constructed with IQ-Tree v2.2.0 (Minh et al., 2020) based on the sequence of *rpoB* genes retrieved from *Bradyrhizobium* genomes (209 new genomes and 566 public genomes downloaded from the NCBI Genbank database). In addition, to identify the diversity of Photosynthetic *Bradyrhizobium* in each sample, a *rpoB* gene tree was also built by combining ASVs and amplified regions from Photosynthetic *Bradyrhizobium* genomes (209 new genomes and 13 public genomes). Although phylogenetic resolution of the *rpoB* gene faded when the short amplified region was used (Fig. S1B), members from each supergroup remain clustered though broken into several subclades, suggesting the use of the tree based on short amplified region has limited effect on amplicon sequence variants (ASVs) assignment using the commonly used phylogenetic placement method (Janssen et al., 2018).

A phylogenomic tree of *Bradyrhizobium* was built using IQ-Tree v2.2.0 (Minh et al., 2020) with our 209 isolates and 566 public genomes (outgroup included) based on 123 shared single-copy genes identified in a previous phylogenomic study of *Bradyrhizobium* (Tao et al., 2021). As our isolates mainly fall into Photosynthetic *Bradyrhizobium* supergroup, we also performed phylogenomic and comparative genomic analyses for this supergroup to understand their phylogenetic and population structure. PopCOGenT (Arevalo et al., 2019) was used to delineate genetically isolated populations for the 222 Photosynthetic *Bradyrhizobium* genomes (13 public genomes and 209 new genomes). The population boundaries delineated by this method is based on recent gene flow barriers, matching well with the idea that bacterial speciation often proceeds rapidly (Arevalo et al., 2019). Next, we selected 28 strains spanning over the main clades and the populations defined by PopCOGenT for symbiotic assays including nodulation and nitrogen fixation capabilities on a tropical legume species *Aeschynomene indica*.

## Results and discussion

### Photosynthetic supergroup of *Bradyrhizobium* is enriched in rice cropland compared to forest and grassland

Both the *rpoB* gene tree (Fig. S1A) and the phylogenomic tree (Fig. S2) grouped *Bradyrhizobium* members into seven supergroups including the Photosynthetic *Bradyrhizobium* (PB). Their largely congruent tree topological structures support the idea that *rpoB* is an appropriate marker gene (Ogier et al., 2019) to distinguish *Bradyrhizobium* supergroups. We therefore designed specific *rpoB* primers to investigate the relative abundance and diversity of each *Bradyrhizobium* supergroup in different terrestrial ecosystems including three plant species (One plant species specific to each ecosystem was chosen, *Oryza sativa* subsp. *Indica* for cropland, *Houttuynia cordata* for grassland, and *Camphora officinarum* for forest). We show that most *Bradyrhizobium* supergroups, including PB, are widely distributed in forest, grassland, and cropland (Fig. 1A, Dataset S1). Among these, PB is notably enriched in the root and rhizosphere niches of rice (Fig. 1B, Dataset S2), implying that members of this supergroup have the potential to be used as plant growth promoting bacteria (PGPB) to promote rice growth and yield.

**Figure 1.**
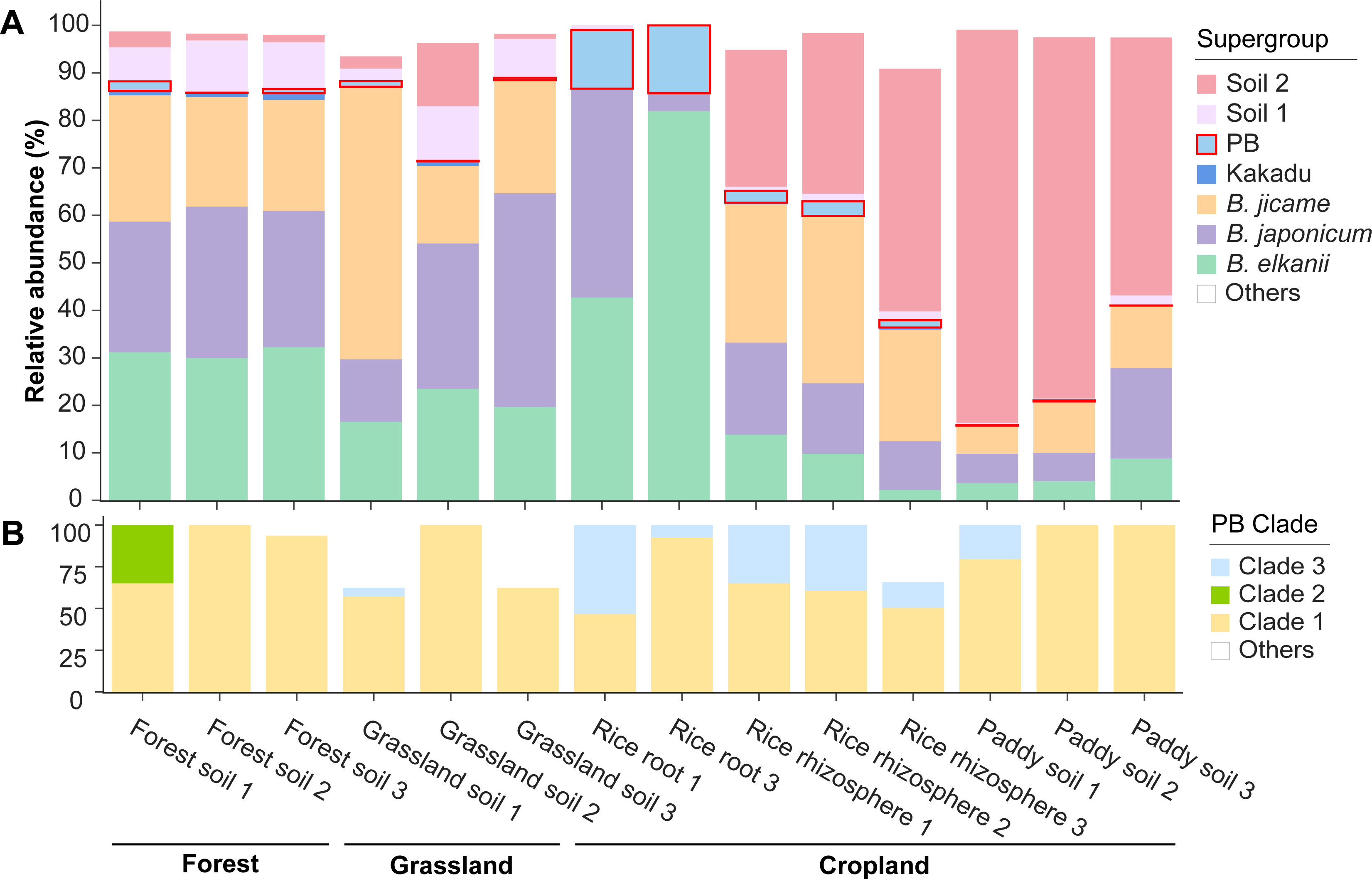
The relative abundance of each *Bradyrhizobium* supergroup (A) and each major clade within the Photosynthetic supergroup of *Bradyrhizobium* (B). The relative abundance of each *Bradyrhizobium* supergroup / PB clade was determined by dividing the number of reads assigned to each supergroup / clade by the total number of filtered reads assigned to *Bradyrhizobium* / PB. Abbreviations: *B. elkanii*, *B. jicamae* and *B. japonicum* represent *Bradyrhizobium elkanii*, *Bradyrhizobium jicamae* and *Bradyrhizobium japonicum* supergroups, respectively.

### Our culture collection contributes a novel deeply branching clade of the Photosynthetic supergroup of *Bradyrhizobium*

With the 209 newly isolated members (Dataset S3), the genome-based phylogeny of the PB supergroup is split into three deeply branching clades (see Fig. S2 for their phylogenetic position in the species tree of the entire *Bradyrhizobium* genus). Clade 1 represents the evolutionarily basal lineage. It initially comprised only 12 strains (Dataset S4), but it now has its diversity expanded by having 68 new strains representing several new lineages delineated as distinct genetically isolated populations (Fig. 2, Dataset S3, S4; also discussed in the next section). Clade 2 has only two strains (the publicly available *Bradyrhizobium* sp. STM3843 and our newly sequenced HKCCYLS1011 isolated from *Oryza sativa* subsp. *japonica*) and represents a phylogenetic intermediate among the three clades. Clade 3 consists of 140 newly isolated strains. The evolutionary branching order of the three clades is verified with two outgroup-independent methods (Fig. 2, Fig. S3) and the outgroup-dependent method (Fig. S2). Collectively, our new strains appreciably increase the existing phylogenetic diversity of the PB supergroup.

**Figure 2.**
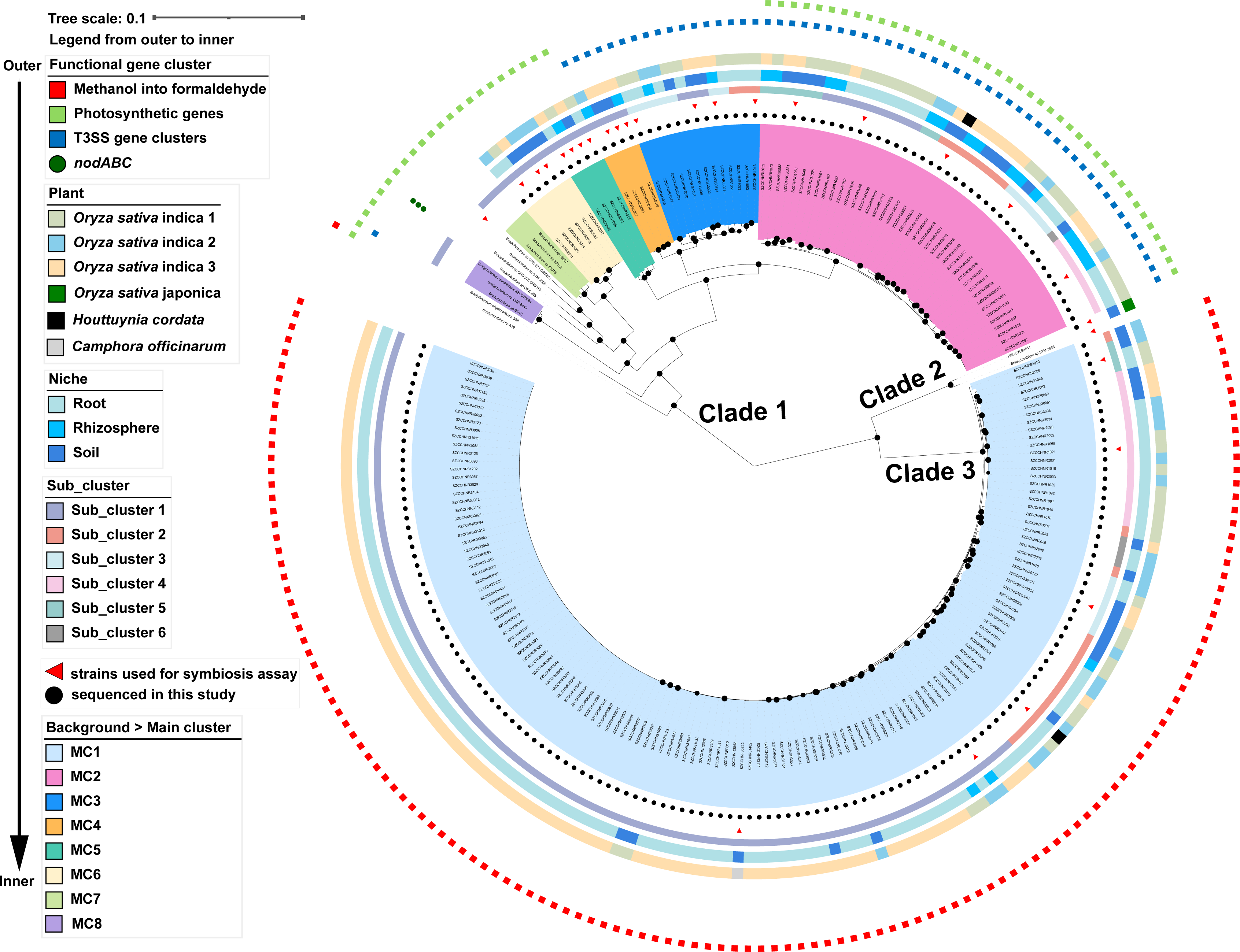
The phylogenomic tree and population delineation of the “photosynthetic” *Bradyrhizobium* (PB) supergroup. The phylogenomic tree was rooted using the minimum variance (MV) method. The 209 PB strains sequenced in the present study were indicated by red dots in the innermost layer surrounding the phylogeny. The ultrafast bootstrap values higher than or equal to 95% calculated by IQ-Tree were labeled on the nodes with black circles. The phylogenomic tree of the Photosynthetic *Bradyrhizobium* based on the minimal ancestor deviation (MAD) rooting method was displayed in Fig. S3.

Comparative genomic analysis shows that except for *Bradyrhizobium* sp. ORS285, which is known to use both a *nod*-dependent and -independent strategy for nodulation (Giraud et al., 2007), all PB supergroup members lack the *nod* genes. Our *rpoB* amplicon analysis showed that PB members in all analyzed ecosystems are dominated by the PB Clade 1, and distribution of the newly isolated Clade 3 is mainly limited in rice-associated niches (Fig. 1B, Dataset S2). It is worth mentioning that the ASVs assigned to Clade 3 are closely related to the cultured members of Clade 3, but none of them have identical sequences to the cultures (Fig. S4). This suggests that future work on cultivation is needed to further expand the fine-scale diversity within Clade 3.

These three major clades may have diverged functionally as each has a unique set of ecologically relevant genes. For example, genes encoding C4-dicarboxylate transporter (*dctBD*) are universally and exclusively found in Clade 1 (Fig. S5, Dataset S5), potentially enabling Clade 1 strains to acquire C4-dicarboxylate compounds (e.g., malate and succinate) from the host plants to fuel the energy-intensive nitrogen fixation (Liu et al., 2018). Other Clade 1-associated genes, though not necessarily exclusively found in Clade 1, include those involved in carbon metabolism and energy conservation processes, such as those encoding histidine utilization enzymes (*hutCFHIU*), glutaconate CoA-transferase (*gctAB*), raffinose/stachyose/melibiose transport system (*msmEFG*), O_2_-independent ubiquinone biosynthesis genes (*ubiDTUV*) and nitrate reductase (*narGHIJ* and *napABCDE*) (Fig. S5, Dataset S5). Specifically, the *hut* pathway is a catabolic pathway that allows using histidine as a source of carbon, nitrogen and energy for growth and to facilitate nitrogen fixation (Bender, 2012). The two non-homologous dissimilatory nitrate reductases (*narGHIJ* and *napABCDE*) and the O_2_-independent ubiquinone biosynthesis genes (*ubiDTUV*) are both necessary for denitrification process (Gregory et al., 2003; Vo et al., 2020).The *narGHIJ* carry out nitrate respiration primarily under anaerobic conditions (Gregory et al., 2003), whereas the *napABCDE* reduce nitrate under both anaerobic and aerobic condition (Gregory et al., 2003).

The genes prevalent to Clade 3 include the *mxa* gene cluster, *mgsABC*, and *mdh12* (Fig. S5, Dataset S5, S6). These genes potentially facilitate the utilization of methanol, which is a waste product during plant-cell wall degradation (Nemecek-Marshall et al., 1995), although a physiological assay did not support this hypothesis (Fig. S6). This suggests that expression of these genes might be regulated by unknown mechanisms. Another unique gene cluster in Clade 3 codes for a protein complex (*exoALOUWY*) (Fig. S5, Dataset S5, S6) probably involved in the synthesis of the major components of exopolysaccharide (EPS), which may induce the immune response of plants (De Sousa et al., 2021; Quelas et al., 2010). Plant LysM kinase receptors perceive EPS, and depending on the compositions, EPS could either impair or promote symbiosis between rhizobia and plants (De Sousa et al., 2021). Other Clade 3-associated genes include malonate decarboxylase (*mdcABCDEG*), UDP-glucose/iron transport system (*STAR12*) and tungstate transport system (*tupABC*) (Fig. S5, Dataset S5). The presence of these genes implies potential unique carbon metabolism (Chohnan and Takamura, 2004) and transporters (Hawkins et al., 2017) in this clade.

Clade 2 consists of only two strains, making it difficult to draw conclusive findings on their gene composition. However, it is still interesting to note that as an intermediate clade, the two strains in Clade 2 uniquely have a gene cluster for the type VI secretion system (T6SS) (Fig. S5, Dataset S5), which is known to influence bacterial competitiveness and symbiosis with eukaryotes (De Sousa et al., 2023; Tighilt et al., 2022). The effects on symbiosis with plants may be closely related to the effectors secreted by T6SS that interact with the host plant and surrounding microbiota (De Sousa et al., 2023). Other specific genes in these two strains include pyruvate ferredoxin oxidoreductase (*porABC*) (Fig. S5, Dataset S5), which exclusively supports fermentation under anaerobic conditions and releases energy at the same time (Wang et al., 2022).

### Fine-scale population structure of the Photosynthetic supergroup of *Bradyrhizobium*

Apart from the broad diversity of the PB supergroup, we further asked whether fine-scale phylogenetic differentiation correlates with the ecological niches where the PB supergroup members were found. As geography may have an important impact on the genetic diversity of rhizobial populations (Greenlon et al., 2019), we performed intensive isolation of PB members from the same rice field to control for the potential impact of geographic separation on PB population structure. Of the 209 new PB supergroup strains, 205 were collected from three rice plants located in the same rice field. Bacterial isolation was performed from within the root, rhizosphere, and bulk soil for each plant, collectively giving rise to 145, 23 and 41 PB strains, respectively (Dataset S3). To facilitate the correlative analysis between fine-scale phylogenetic groups and the niches, we assigned isolates’ genomes into populations defined by PopCOGenT (reported as "main cluster" or MC) (Arevalo et al., 2019). As bacterial members within a population have significantly higher recombination rate than those across populations, “population” defined here aligns with the “species” in higher eukaryotes. Additionally, PopCOGenT can detect subpopulations (reported as ’subclusters’), which are under ongoing differentiation within a population.

Using this approach, we show that i) all strains in Clade 3 share the membership of a single population (MC1), ii) the two strains in Clade 2 each form a distinct population (MC id not given), though they may be deemed as members of a single operational species as their genome-wide average nucleotide identity (ANI) is 96.5% (Fig. S7, Dataset S7), exceeding the species threshold of 95%, and iii) strains from Clade 1 fall into numerous populations, among which, MC2, MC3, MC4, MC5 and MC6 are each exclusively comprised of the new strains, whereas MC7, MC8 and the remaining six unassigned isolates each forming a distinct population are all publicly available (Fig. 2). In Clade 1, the within- and between-population similarity is above and below 95% ANI (Fig. S7, Dataset S7), respectively, consistent with the operational species threshold of 95% ANI (Konstantinidis and Tiedje, 2005), whereas the 16S rRNA gene shows little divergence with the between-population similarity generally above 99% (Fig. S7, Dataset S7). Based on the available samples, intra-population subdivision occurred within MC1, MC2 and MC3. We found that each population and subpopulation have members sampled from different plants and niches (Fig. 2), suggesting that the PB populations sampled from the rice filed are not genetically subdivided according to the physical separation between the plant individuals or following the niche separation between root, rhizosphere, and bulk soil.

We also found interesting associations between important metabolic pathways and population identity. Remarkably, although the PB supergroup was named by the presence of the photosynthetic genes (Avontuur et al., 2019), these genes are exclusively and universally found in MC2, MC5, MC6, MC7, MC8 and the unassigned individuals in Clade 1 but completely missing from MC3 and MC4 of Clade 1, Clade 2 and Clade 3 (MC1) (Fig. 2, Dataset S6). Our result indicates that photosynthesis is not a characteristic trait that defines the PB supergroup. Also interesting is the prevalence of a unique Type III secretion system (T3SS) in MC2, MC3 and MC4 but completely missing in other populations (Fig. 2, Fig. S8B, S8C, Dataset S6). It is one of the six T3SS subtypes identified from known *Bradyrhizobium* phylotypes (Teulet et al., 2022; Teulet et al., 2020). It was previously identified only in the PB strain *Bradyrhizobium oligotrophicum* S58 (thus named ‘S58-T3SS’ subtype) and its function remains unknown (Teulet et al., 2020). Since the general function of T3SS is to translocate effector proteins into host cells that modulate the host immune response (Teulet et al., 2022), it cannot be excluded that this T3SS type specifically identified in PB members plays an important role during their interaction with their host plants (*Aeschynomene* spp. and rice).

### The ability to nodulate *Aeschynomene indica* is a conserved trait shared by Photosynthetic supergroup of *Bradyrhizobium* but differs between the major clades

All PB supergroup members have a *nif* gene cluster necessary for nitrogen fixation. Duplication of the *nifH* gene which encodes one of the structural proteins of the nitrogenase enzyme complex was observed in all PB strains (Fig. S8A), but the functional consequence is not known. Across the PB supergroup members, an important difference was observed regarding how the *nif* genes are structured. In most populations, all *nif* genes are co-located, along with other genes (e.g., *sufBCDX*, *glbO* and *fixABCX*) potentially involved in nitrogen fixation. This is not the case for a few members such as some basal lineages (e.g., *Bradyrhizobium* sp LMG 8443) and some members assigned to several populations (e.g., SZCCHNR1015 in MC5 and *Bradyrhizobium* sp. 83002 in MC7), where the *nif* genes are present in two contigs. This is likely due to structural rearrangement, but a DNA assembly artefact cannot be ruled out.

A common feature observed of the PB *nif* cluster is the universal presence of the *nifV* gene encoding for a homocitrate synthase that is involved in the biosynthesis of the nitrogenase cofactor (FeMo-Co), which is absent in most other rhizobial lineages (Hakoyama et al., 2009; Nouwen et al., 2017). An acetylene reduction assay on 28 representative PB strains, which cover all three major clades, all newly identified populations (MC1 to MC6), most of the identified subpopulations within each population, and the model PB strain ORS278 as control, confirmed their ability to fix nitrogen under free-living conditions (Fig. S9A), a trait absent in most *nod*-carrying rhizobia (Nouwen et al., 2017).

The plant inoculation experiments showed that all 28 representative strains were also able to induce nodules on *A. indica* (Fig. 3B, 3C). However, a significant difference in symbiosis efficiency was observed between the major clades. Clade 1 strains had the same symbiotic properties as the model strain ORS278, inducing many nitrogen-fixing nodules that stimulated the growth of the plants (Fig. 3A, 3B, Fig. S9B, S10A). In contrast, Clade 3 strains formed much fewer nodules compared to Clade 1 strains and the measured nitrogenase enzyme activity was lower under the symbiotic condition (Fig. 3B, Fig. S9B). Microscopic analysis showed that the nodules elicited by Clade 3 strains displayed multiple aberrant phenotypes: i) some nodules contained necrotic areas, ii) in others the central tissue was completely digested, and finally, iii) the nodules that displayed less drastic symptoms contained mainly dead bacteria (Fig. 3C and 3F). These observations indicate that besides inducing fewer nodules, Clade 3 strains are unable to maintain an effective chronic infection, which explains why they have no beneficial effect on plant growth (Fig. 3A, 3C, 3F). Clade 2 represented by only two strains STM3843 and HKCCYLS1011 exhibits a transitional pattern of phenotype, consistent with its intermediate phylogenetic position between Clade 1 and Clade 3. The number (Fig. 3B) and phenotype (Fig. 3C, 3E) of the nodules elicited by strain STM3843 are comparable to those of Clade 1 strains (Fig. 3B, 3C, 3D) though stimulation of plant growth is less (Fig. 3A), whereas the nodules stimulated by strain HKCCYLS1011 (Fig. 3B, 3C, 3E) are like those of Clade 3 strains (Fig. 3B, 3C, 3F) and no effect on plant growth was observed (Fig. 3A).

**Figure 3.**
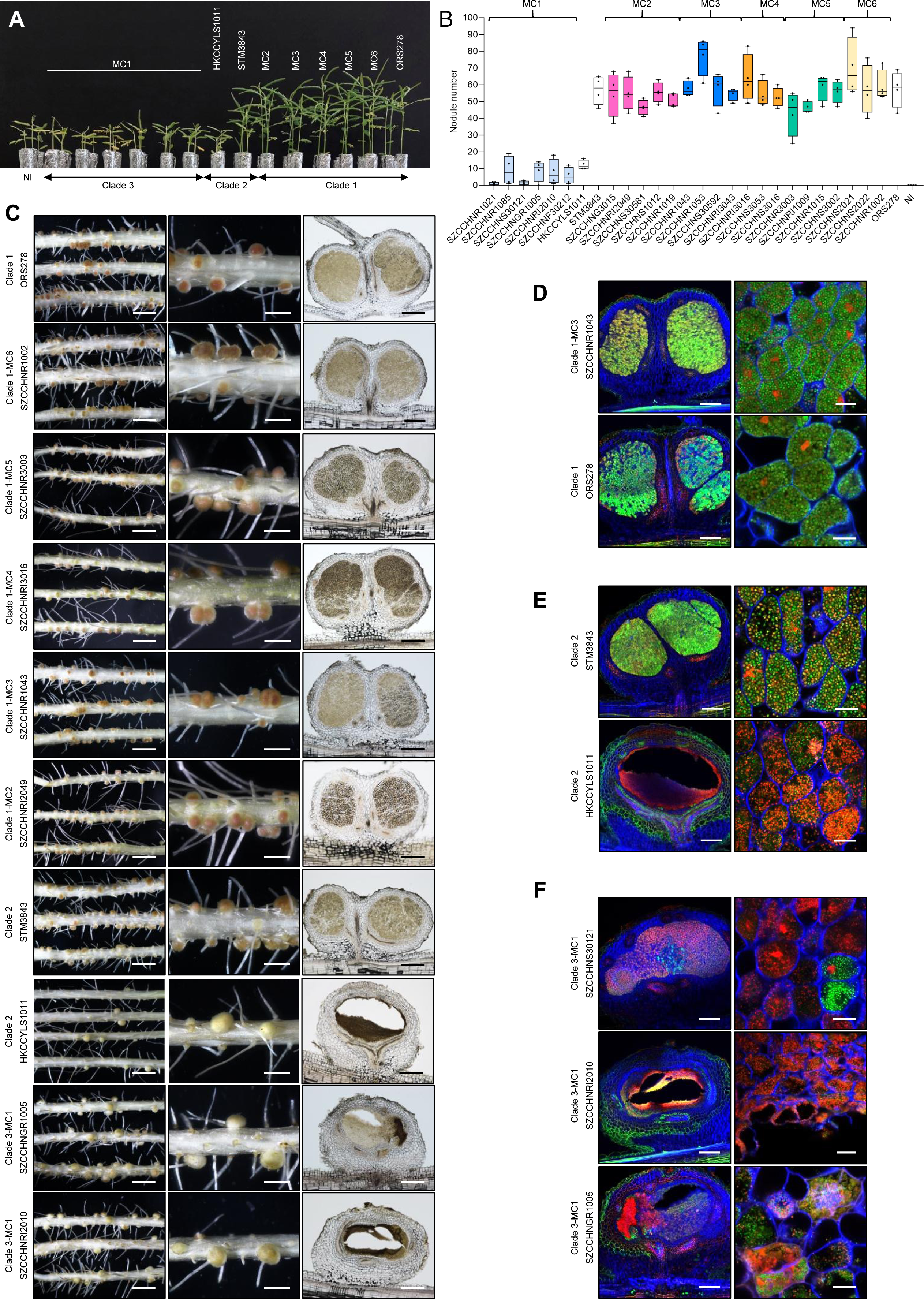
Symbiotic properties of several representative strains of the main populations identified in the PB supergroup. (A) Comparison of growth of the *A. indica* plants (leaf phenotype) non-inoculated (NI) or inoculated with different representative strains of PB. The six MC1 representative strains are presented in the following order: SZCCHNR1021, SZCCHNR1085, SZCCHNS30121, SZCCHNGR1005, SZCCHNRI2010 and SZCCHNF30212. For the other MCs, only one representative strain is shown: SZCCHNRI2049 (MC2), SZCCHNS30592 (MC3), SZCCHNRI3016 (MC4), SZCCHNS3002 (MC5) and SZCCHNS2021 (MC6), Photos were taken 17 days after inoculation. (B) Nodule number on *A. indica* plants induced by different representative strains of PB at 17 days after inoculation. Box plots show the results of one of the two experiments performed independently (4 plants each). (C) Symbiotic phenotypes of some representative strains tested on *A. indica*. Column 1 and 2: photo of roots and nodules. Scale bars: column 1, 0.5 cm; column 2, 0.2 cm. Column 3: Micro-sections of nodules observed using light microscopy. Scale bars: 250 µm. (D) Confocal microscopy images of micro-sections of nodules elicited by Clade 1 strains. The nodules formed by SZCCHNR1043 from MC3 and ORS278 are shown as examples. The central nodule tissue is intracellularly infected with live spherical bacteroids stained green by SYTO9. (E) Confocal microscopy images of micro-section of nodules elicited by Clade 2 strains (STM3843 and HKCCYLS1011). The nodules elicited by STM3843 look normal but a mix of live and dead bacteria (stained red by propidium iodide) can be noticed while the nodules formed by HKCCYLS1011 displayed a central infected tissue that is digested. (F) Confocal microscopy images of micro-section of nodules elicited by Clade 3 strains displaying different phenotypes: nodule containing mainly dead bacteria (SZCCHNS30121), nodule with a completely digested central tissue (SZCCHNRI2010); and nodule with a necrotic area and a digested zone (SZCCHNGR1005). Scale bars for (D), (E) and (F): column 1, 200 µm; column 2, 10 µm.

The symbiosis efficiency differences observed between the major clades may result from an inability of the Clade 3 strains to cope with the plant’s immune response, and/or an over-induction of the plant’s defense mechanisms due to the absence or not properly recognized symbiotic signal(s). However, it is remarkable that all strains from the PB supergroup, which come from different origins and geographical locations, are capable of nodulating *A. indica*. The ability to develop an NF-independent nodulation with *Aeschynomene* spp. thus appears to be a common trait of PB supergroup members. Thus, unlike the *nod* genes of other rhizobial lineages which are accessory genes acquired by horizontal transfer, the genes governing NF-independent symbiosis may be essential genes belonging to the core genome of the PB supergroup.

## Concluding remarks

The results of *rpoB* amplicon sequencing of samples from forest, grassland, and rice field indicate that members of the Photosynthetic *Bradyrhizobium* (PB) are enriched in rice field. By large-scale isolation and genome sequencing, we report a novel major clade, thereby greatly expanding the phylogenetic diversity of the cultured members of the PB supergroup. An important finding is that although all assayed phylogenetically diverse PB strains can nodulate *Aeschynomene* spp., the branching order of the three major clades correlates nicely with their symbiosis efficiency, with the clade taking the intermediate phylogenetic position showing the symbiosis efficiency in between. It is therefore likely that the ability to establish symbiosis with *Aeschynomene* plants, which often grow in the same wetlands as rice, is a key factor that shapes the broad diversity of the PB supergroup. At the population level, no significant difference in symbiosis efficiency was observed, suggesting that other traits such as photosynthesis and T3SS that are uniquely associated with some but not all populations, might be among the important drivers of population-level genetic differentiation and ecological adaptation. Collectively, our study provides insights into the ecology and evolution of the Photosynthetic supergroup within the globally dominant soil bacteria *Bradyrhizobium*, and additionally serves as a prime example that links deep phylogenetic diversity of an ecologically relevant bacterial group to their major phenotypic and functional diversity.

## Acknowledgements

We thank Xiaoyuan Feng for helping with genome assembly. This work was supported by the Hong Kong Research Grants Council Area of Excellence Scheme (AoE/M-403/16) and a grant from the French National Research Agency (“ET-Nod”; ANR-20-CE20-0012).

## Data availability

The genomic sequences and raw reads of the 209 newly sequenced *Bradyrhizobium* isolates have been uploaded to NCBI (Project ID: PRJNA983111).

## Supplementary Information

### This PDF file includes

Supplementary Text. Methods

Supplementary references.

Figures S1 to S11.

## Supplementary Text: Methods

### Sample niche classification

Excess soil was vigorously shaken away from the plant roots to remove loose soil, leaving approximately 1 mm of soil attached to the roots, which constitutes the rhizosphere compartment (Edwards et al., 2015). To separate the 1 mm of rhizosphere soil directly from the roots, roots were cut into 5 cm lengths and two grams were placed in a sterile flask containing 50 mL of sterile phosphate buffered saline (PBS) solution and vortexed several times (15 seconds). The PBS solution used to wash the roots was transferred to a 50 mL Falcon tube and then centrifuged at 10,000 x g for one minute. The supernatant was discarded, and the remaining soil samples were saved as rhizosphere samples (Edwards et al., 2015). The roots were then resuspended and vortexed three times (15 seconds each) before being transferred to new Falcon tubes as root samples. Soil away from the root was stored as bulk soil.

### Bacterial isolation

The root, rhizosphere, and bulk soil samples were placed in the incubation chamber for three to four days at 28°C and 70% relative humidity under the day/night (16/8 h) cycle. All root samples were washed with sterile Milli-Q water to remove any adhering soil, then subjected to the surface sterilization by immersion in 75% ethanol for three minutes, followed by 5% sodium hypochlorite for five minutes, and rinsed five times with sterile distilled water (Mbai et al., 2013). The washed roots were then aseptically dried on the clean bench before being ground in a sterilized mortar and pestle with PBS buffer (pH 6.8). A series of fivefold and tenfold dilutions were prepared and then inoculated (100 µL) on modified arabinose-gluconate (MAG) media without any nitrogen source (1.0 g DL-arabinose, 1.0 g sodium gluconate, 1.0 g yeast extract, 2 mL KH_2_PO_4_ solution (110 g/L), 4 mL Na_2_SO_4_ solution (62.5 g/L), 1 mL MgSO_4_•7H_2_O solution (180 g/L), 1 mL CaCl_2_ solution (13 g/L), 1 mL FeCl_3_•6H_2_O solution (6.7 g/L) and 15 g agar were added. The medium was made up to 1 L with Milli-Q water, the pH was adjusted to 6.6 with KOH, and then autoclaved at 121 ℃ for 15 - 30 minutes.) (Tao et al., 2021). To prepare soil inoculations, 5.0 g of fresh soil was mixed with 45 mL of sterile deionized water in a 50 mL Falcon tube. One mL of soil suspension was serially diluted 10-fold after mixing with a vortex mixer, and 100 μL of diluted samples were spread on the MAG media (Tao et al., 2021). The isolation plates were then placed in a 28 °C incubator to allow bacteria to grow. After one week, colonies with specific morphology (small colonies with white or pink color) were picked and purified on fresh MAG media with the addition of 2 mL NH_4_Cl solution (160 g/L). Purified strains were preserved at -80 °C in glycerol suspensions (30% v/v).

To identify the taxonomy of each strain, colony polymerase chain reaction (PCR) was performed. Briefly, bacterial colonies were mixed with the 10% (w/v) Chelex solution [Bio-Rad, USA] and incubated at 90 °C for 20 minutes to release DNA from bacterial colonies as templates. The universal bacteria 16S rRNA primers 27F (5′-AGRGTTYGATYMTGGCTCAG-3′) and 1492R (5′-GGYTACCTTGTTACGACTT-3′), Premix Taq [Takara Bio, USA] and RNA/DNAse- free water were used in the PCR. The PCR condition included an initial denaturation at 95 °C for five minutes, followed by 35 cycles of amplification (95 °C for 45 s, 55 °C for 45 s and 72 °C for 90 s) and a final extension at 72 °C for 10 minutes (Tao et al., 2021). PCR products were run on 0.8% agarose and viewed by the automatic analysis system of electrophoresis gel imaging. Positive PCR products were sequenced in BGI Genomics. The taxonomic information of each isolate was analyzed by comparing the 16S rRNA gene sequences with *Bradyrhizobium* in EzBioCloud (Yoon et al., 2017).

### Primer design for detecting *Bradyrhizobium* diversity

It is well known that the closely related genera are difficult to be distinguished by the traditional 16S rRNA gene sequences due to their slow evolutionary rate (Vos et al., 2012). In contrast, the *rpoB* gene has a higher resolution and can be used as a marker gene to distinguish closely related species (Vos et al., 2012). Therefore, we designed specific *Bradyrhizobium rpoB* primers for amplicon sequencing to investigate the relative abundance and diversity of *Bradyrhizobium* supergroups in different samples. The primer sequences were BR2106F (CCGRTSACGCCBGACAAG) and BR2516R (TGTCGCCCTTCYTGACGAYR), producing a ∼410 bp sequence. The primers were validated to amplify various *Bradyrhizobium* members as well as strains within the PB supergroup with specificity (Fig. S11). PCR conditions included denaturation at 95 °C for three minutes, followed by 35 cycles of 95 °C for 30 seconds, 61 °C for 45 seconds and 72 °C for 45 seconds, with a final extension at 72 °C for 10 minutes.

### Amplicon sequence analysis

The quality and quantity of the DNA extracted from the soil and root samples were assessed using a NanoDropTM 2000 [Thermo Fisher, USA]. The amplicon sequence variants (ASVs) obtained from the *rpoB* amplicon sequences after denoising were subsequently filtered out the non-*rpoB* amplicon by conducting BLASTN searches against the NCBI nr database. The aligned ASVs with an e-value lower than 1e-10 but without alignment with ‘DNA-directed RNA polymerase subunit beta’ in the nr database were removed.The reference *rpoB* sequence was extracted from the genomic sequence and then aligned. The alignment was trimmed to retain the amplified region and used as the reference alignment. The ASVs were then aligned with the reference alignment using PaPaRa v2.5 (Berger and Stamatakis, 2012). The aligned ASVs were subsequently placed on the phylogenetic tree (Fig. S1B) using EPA-ng v0.3.8 (Barbera et al., 2019) and assigned to seven *Bradyrhizobium* supergroups, including Soil 1, Soil 2, *Bradyrhizobium elkanii*, *Bradyrhizobium jicamae*, Kakadu, Photosynthetic and *Bradyrhizobium japonicum* supergroups. The relative abundance of each *Bradyrhizobium* supergroup was determined by dividing the number of reads assigned to each *Bradyrhizobium* supergroup by the total number of filtered reads assigned to *Bradyrhizobium* (Fig. 1a, Dataset S1). Similarly, the relative abundance of each clade of Photosynthetic *Bradyrhizobium* (PB) was calculated by dividing the number of reads assigned to each PB clade by the total number of filtered reads assigned to PB (Fig. 1b, Fig. S3, Dataset S2).

### Basic soil characteristics

Soil samples were separated to measure the basic soil characteristics after all the samples were brought back to the laboratory. Soil water content was determined by drying fresh soil (10.00 g) at 105°C for 6 hours. Soil pH was measured at the ratio of 1:5 (w/w) of soil-to-deionized water using a pH electrode. The potassium dichromate oxidation process combined with the heating method was used to measure the soil organic carbon (SOC). Total nitrogen (TN) was determined by the kjeldah method (Bao, 2000). Total phosphorus (TP) was identified based on the sodium hydroxide melting-molybdenum antimony colorimetric method (Bao, 2000).

Available phosphorus (AP) was ascertained using the molybdenum antimony anti-colorimetric method (Bao, 2000). Soil nitrite nitrogen (NO_2_^-^), nitrate nitrogen (NO_3_^-^) and ammonium nitrogen (NH_4_^+^) were extracted with KCl solution (2 mol L^-1^) and detected by continuous flow analysis (Tian et al., 2014). Dissolved organic carbon and nitrogen (DOC and DON) were analyzed using a TOC/TN analyzer (Shimadzu, Analytical Sciences, Kyoto, Japan). Microbial biomass carbon and nitrogen (MBC and MBN) were determined by chloroform fumigation extraction (Durenkamp et al., 2010; Wu et al., 1990). Fe and Mo contents were measured by using inductively coupled plasma mass spectrometry (ICP-MS). These sample-associated metadata were provided in Dataset S3.

### Genome sequencing, assembly and annotation

Bacteria Genomic DNA Extraction Kit [OMEGA Bacterial DNA Kit D3350] was used to extract genomic DNA from each of the 209 isolates. NanoDrop^TM^ 2000 [Thermo Fisher, USA] was used to assess the quality of the extracted DNA samples with the following criteria, A260/A280 > 1.8, A260/A230 > 2.0 and A260/A270 > 1.0. Whole genome sequencing was performed in Wuhan Huada Gene Biotechnology Company by using the MGISEQ-2000 PE150+150+10+10 platform (paired-end reads of 150 bp). The BBMerge method in the BBmap package v38.79 was used to identify the untrimmed adapters associated with the raw reads (Bushnell et al., 2017). Subsequently, Trimmomatic v0.39 was used to trim adapters and low-quality reads (Bolger et al., 2014). Reads with less than 40 bp in length were discarded, and the quality of the remaining reads was assessed by FastQC (https://www.bioinformatics.babraham.ac.uk/projects/fastqc/). Contigs were assembled using SPAdes v3.10.1 based on the remaining high-quality paired-end reads with default parameters (Bankevich et al., 2012). Those contigs with lengths greater than 1,000 bp and a k-mer coverage greater than five were kept for further analysis. The quality of the genome assemblies (Dataset S8) was assessed by CheckM v1.0.7 (Parks et al., 2015). In general, 68 newly sequenced strains were clustered with 12 publicly available strains as Clade 1, strains STM3843 and the newly sequenced HKCCYLS1011 were clustered as Clade 2, and the remaining 140 newly sequenced strains were clustered as Clade 3. Genomes from Clade 3 exhibited a significantly lower GC content (63.67 ±0.07% vs. 65.39 ±0.22%, *p* < 0.001, phylogenetic ANOVA) and a similar genome size (7.45 ±0.14 Mb vs. 7.56 ±0.33 Mb, *p* = 0.746, phylogenetic ANOVA) compared to Clade 1 (Datasets S2 and S7).

We noted an assembly error related to the *nif* island that was supposed to harbor two *nifH* genes. This was due to the presence of two *nifH* gene copies with nearly identical sequences, which can lead to mis-assembly in short-read sequencing technologies such as Illumina or MGISEQ-2000 (Tørresen et al., 2019). To address this issue, we mapped all raw reads to the assembled genome and observed that the sequencing depth of the *nifH* gene was twice that of other nearby genes. We further conducted Nanopore sequencing on three phylogenetically distantly related PB strains (SZCCHNR3119, SZCCHNS2021 and SZCCHNS1050), and had their genomes assembled using Flye v2.6 (Kolmogorov et al., 2019). Analysis of these genomes confirmed the presence of two *nifH* genes separated by approximately 43 genes in these strains.

To correct this error in the remaining PB strains, we re-assembled their genomes using the reference-guided method (Cabuk and Unlu, 2022) in Spades v3.10.1 with the parameter of “-- untrusted-contigs” (Bankevich et al., 2012). For each assembled genome, genes were predicted using Prokka v1.12 (Seemann, 2014) and annotated using the Cluster of Orthologous Genes (COG) database (Galperin et al., 2020) and the Kyoto Encyclopedia of Genes and Genomes (KEGG) database (Kanehisa et al., 2023; Kanehisa et al., 2004) .

### Phylogenomic tree construction for all available *Bradyrhizobium* lineages

A phylogenomic tree for all available *Bradyrhizobium* lineages (209 new genomes and 566 public genomes downloaded from the NCBI Genbank database) was constructed by IQ-TREE v2.2.0 (Minh et al., 2020). The parameter “-s alignment -spp partition -m MFP -mset LG,WAG,JTT -mrate E,G,I,G+I -bb 1000” was applied so that each gene was allowed to have its own best-fit substitution model, automatically selected by the ModelFinder implemented in IQ-TREE v2.2.0 (Minh et al., 2020). The branch support was assessed by 1000 ultrafast bootstrap approximations (Hoang et al., 2017).

### Phylogenomic and comparative genomic analyses for Photosynthetic *Bradyrhizobium*

OrthoFinder v2.3.4 (Emms and Kelly, 2019) was used to identify orthologous gene families for all Photosynthetic *Bradyrhizobium* strains. Each of the identified 3,300 single-copy ortholog families was aligned at the amino acid level using MAFFT v7.487 (Katoh and Standley, 2013), and trimmed using trimAl v1.4.rev15 with the parameters “-automated1 -resoverlap 0.55 - seqoverlap 60” (Capella-Gutiérrez et al., 2009). A maximum likelihood phylogenomic tree of the PB supergroup was built based on the concatenated alignment of 3,300 single-copy orthologs using IQ-TREE v2.2.0 with 1,000 ultrafast bootstrap replicates (Hoang et al., 2017). The best-fit evolutionary model for each ortholog was determined by ModelFinder implemented in IQ-TREE v2.2.0 (Minh et al., 2020).

As shown in Fig. S2, the Kakadu supergroup was considered to be a sister lineage to the Photosynthetic supergroup. However, the long branches connecting these two supergroups suggested that using the Kakadu supergroup as an outgroup to root the species tree of the Photosynthetic supergroup may not be appropriate. Therefore, the Photosynthetic supergroup phylogenomic tree was rooted using outgroup-independent rooting methods, specifically the minimum variance (MV) (Mai et al., 2017) and the minimal ancestor deviation (MAD) (Tria et al., 2017) methods. The MV method identifies the root with the minimum variance of root-to-tip distances, while the MAD method considers all branches as plausible root positions, calculates the relative deviation from the clock-likeness for each candidate, and determines the root with the minimal mean relative deviation from the molecular clock interpretation of all branches (Tria et al., 2017). The trees based on these two outgroup-independent methods indicated the same root position (Fig. 2 and Fig. S3). To compare continuous traits of strains while incorporating the evolution of those traits, a phylogenetic ANOVA (Garland et al., 1993) was conducted using the ‘phylANOVA’ function with 1,000 simulations implemented in the R package ‘phytools’ (Revell, 2023).

The similarity of the PB strains was measured at both 16S rRNA gene and whole-genome levels. For the former, pairwise 16S rRNA gene identity was calculated using BLAST (Boratyn et al., 2013), and strains were clustered using the complete linkage method based on sequence identity. For the latter, the whole-genome average nucleotide identity was estimated by FastANI v1.2 (Jain et al., 2018).

### Symbiotic analysis on *Aeschynomene indica*

The 28 strains used for symbiotic analysis were grown in AG medium (Sadowsky et al., 1987) at 28°C on Petri dishes for one week. Bacteria were harvested from the plate and resuspended in 10 mL of sterile water and the OD_600_ was adjusted to 1.0. *A. indica* plants were cultured as previously described (Okazaki et al., 2016). Each strain was inoculated into four plants with 1 mL of bacterial suspension and the symbiotic properties (number of nodules per plant and nitrogenase enzyme activity) were analyzed at 17 days after inoculation (Bonaldi et al., 2010). Cytological analysis of nodules elicited by one strain representative of the populations (MC2 to MC6), two strains for the Clade 2 and three strains for the MC1 population was performed as described (Songwattana et al., 2021). Each strain was assessed in duplicate.

### In vitro nitrogenase enzyme activity

Bacteria were grown in 9 mL vacuette® tubes (Greiner Bio-One GmbH) containing 2 mL of 0.8% agar BNM-B medium with 10 mM succinate and 10 mM arabinose as carbon source at 28 °C (Nouwen et al., 2017). At the beginning of the experiment, 10% acetylene was added to the vacuette® tubes and after eight days of incubation, the amount of ethylene produced by the bacterial culture was measured by gas chromatography (Giraud et al., 2000).

### Methanol metabolism

We selected 17 strains covering the major populations predicted by PopCOGenT. The reference strain *Bradyrhizobium* diazoefficiens USDA110 was used as a positive control. Each strain was inoculated into 14 mL test tubes containing 5 mL minimal salts medium, supplemented with 30 μM CeCl_2_ (lanthanide chlorides) and 0.5% methanol as the sole carbon source (Wang et al., 2019) and incubated at 28 ℃ with reciprocal shaking at 280 rpm. Growth was monitored spectrophotometrically by measuring the optical density at 600 nm (OD_600_).

---

**Figure S1.**
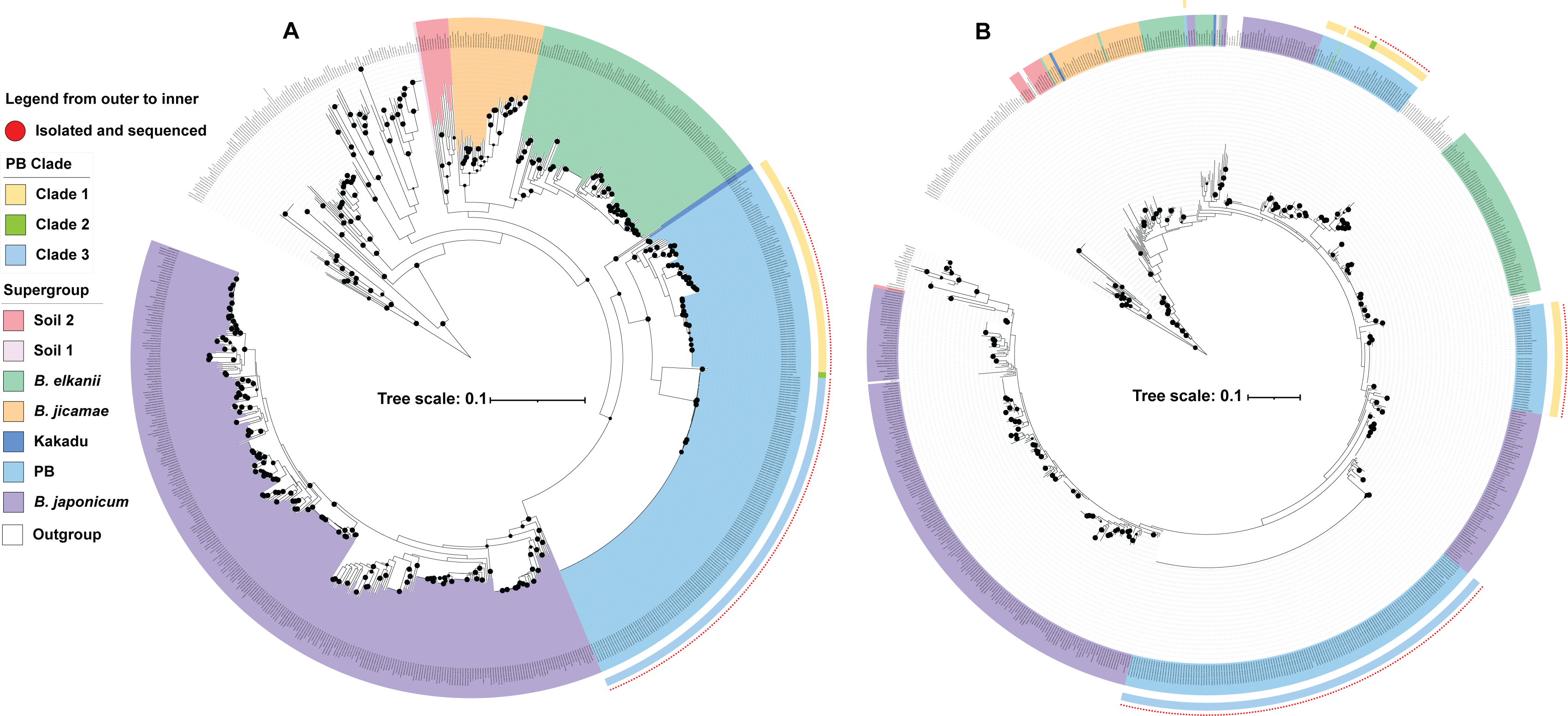
*RpoB* gene tree constructed by full length (A) and amplified region (B). Strains from Xanthobacteraceae were used as an outgroup. The black circles on the nodes indicate ultrafast bootstrap values higher than or equal to 95% calculated by IQ-Tree. The 209 PB strains sequenced in the present study are indicated by red dots in the outermost layer surrounding the tree.

**Figure S2.**
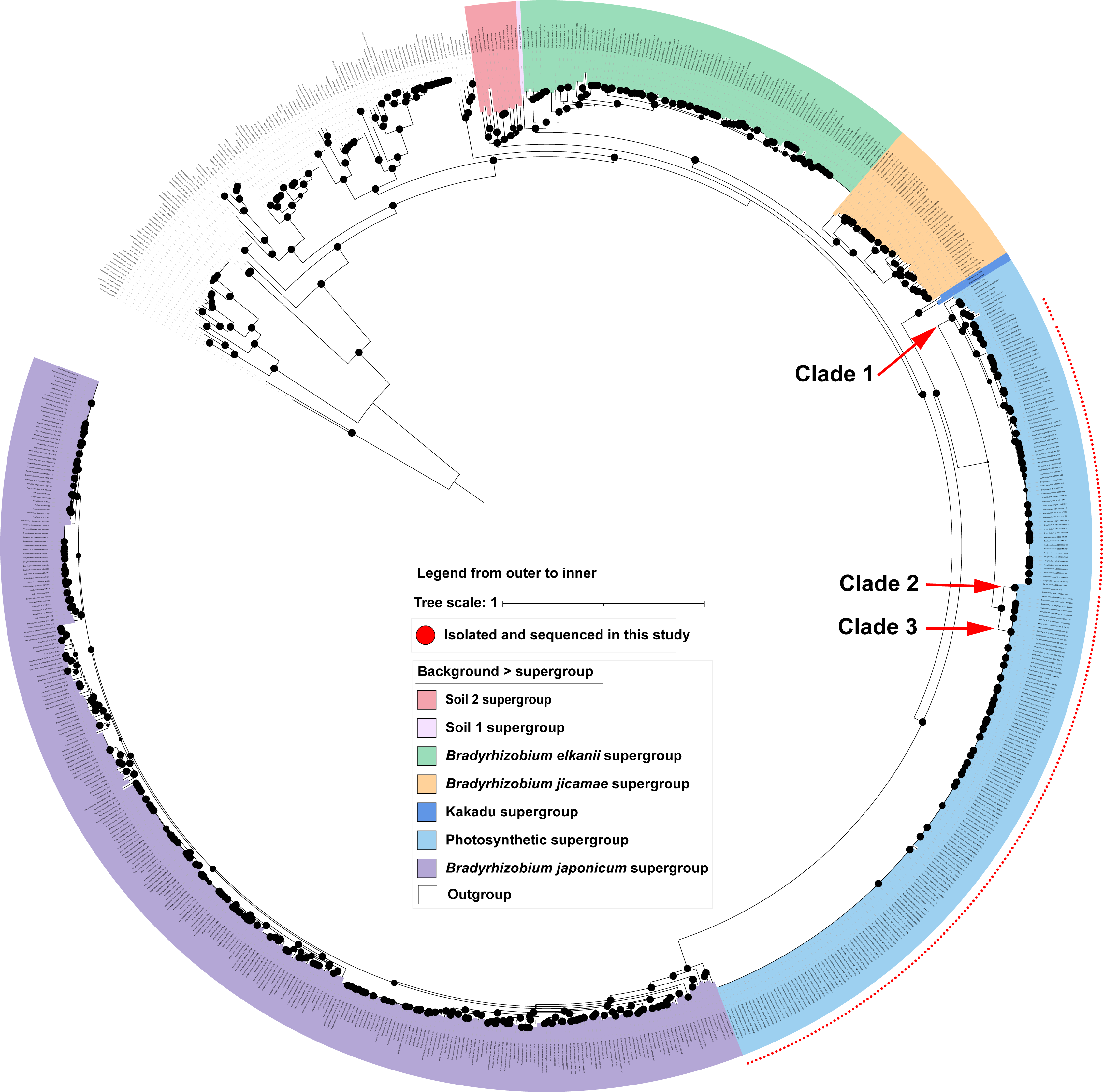
The maximum-likelihood phylogenomic tree of *Bradyrhizobium*. Strains from Xantho-raceae were used as an outgroup. The tree was constructed using the 123 orthologous genes. fied in a previous study (Tao et al., 2021). The black circles on the nodes indicate ultrafast trap values higher than or equal to 95% calculated by IQ-Tree. The 209 PB strains sequenced present study are indicated by red dots in the outermost layer surrounding the tree.

**Figure S3.**
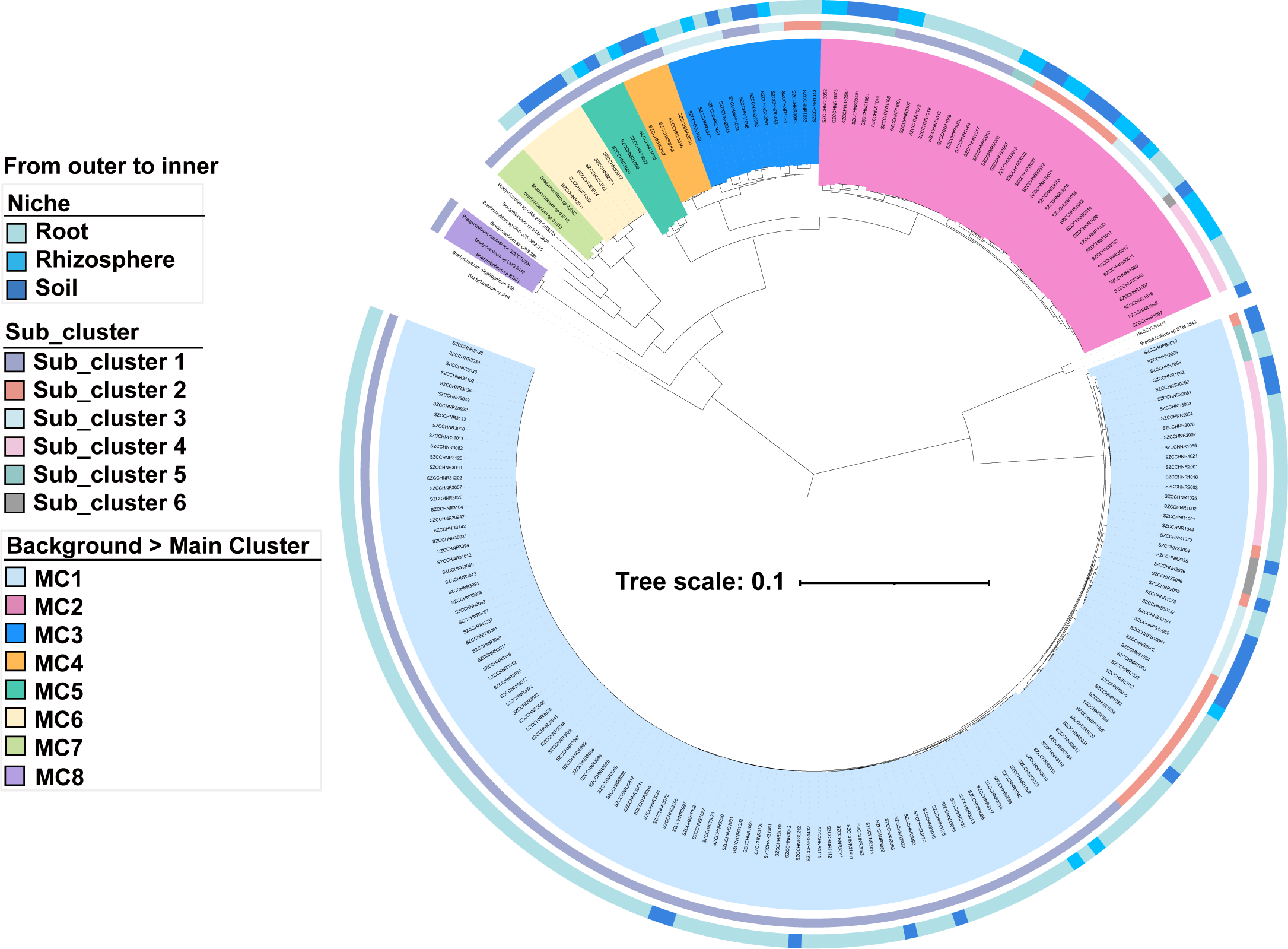
The phylogenomic tree of the photosynthetic *Bradyrhizobium* is based on the minimal ancestor deviation (MAD) rooting method. Solid circles in the phylogeny indicate nodes with IQ-Tree’s ultrafast bootstrap values ≥ 95%.

**Figure S4.**
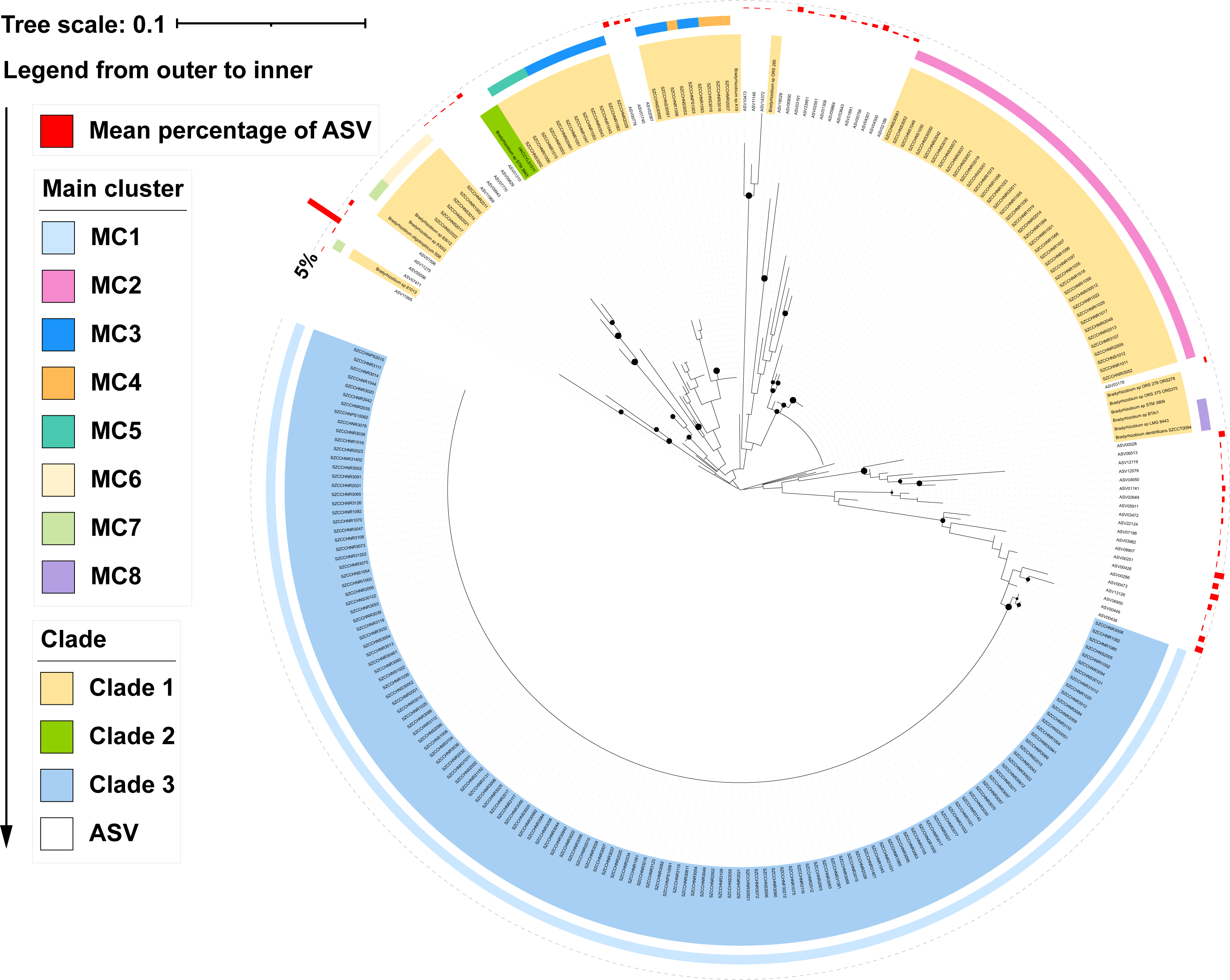
The ASVs (amplicon sequence variants) and *rpoB* genes (amplified region from photosynthetic *Bradyrhizobium* genomes) tree. This gene tree was rooted by the minimum variance (MV) method. The 222 *rpoB* genes from PB genomes in this study were divided into each clade and main cluster (MC) according to Fig. 2. The ultrafast bootstrap values higher than or equal to 95% calculated by IQ-Tree were labeled on the nodes with black circles.

**Figure S5.**
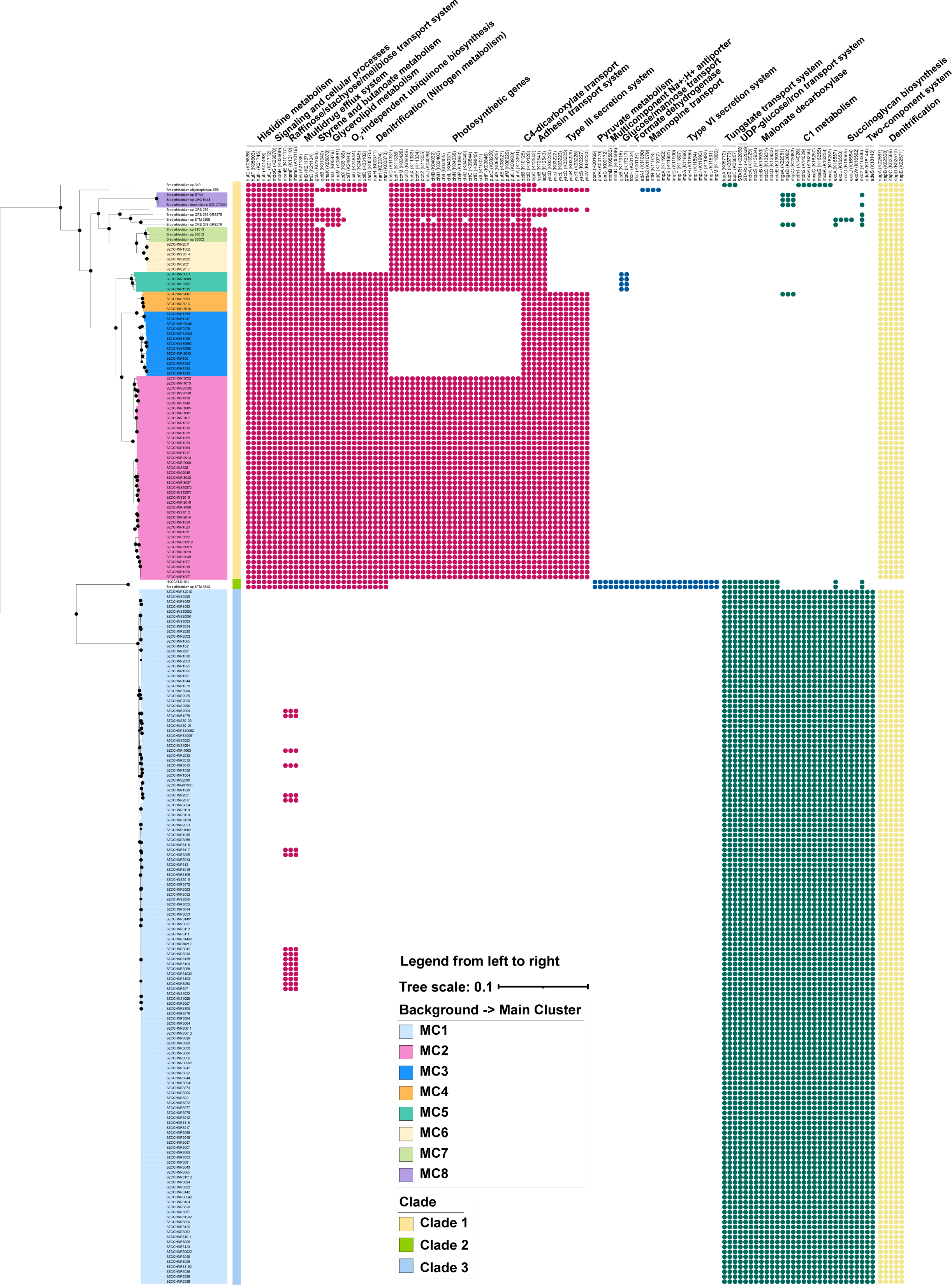
The phyletic pattern of specific genes between three PB clades. The solid circles and blank in the right panel represent the presence and absence of the genes, respectively. The genome tree in the left panel was displayed in rectangular mode. Detailed information of specific genes was shown in Dataset S5. Histidine metabolism *hutCFHIU*, signaling and cellular processes *mdoGH*, raffinose/stachyose/melibiose transport system *msmEFG*, multidrug efflux system *triABC*, styrene and butanoate metabolism *gctAB*, glycerolipid metabolism *dhaKLM*, O_2_-independent ubiquinone biosynthesis *ubiXTUVD*, nitrogen metabolism (denitrification) *narGHIJ 5napABCDE*, photosynthetic genes 1) porphyrin metabolism *bchCFMOXYZJ* and *chlBHILNPG* 2) carotenoid biosynthesis *crtCDIF* 3) light-harvesting complex *pufAB* 4) photosynthetic reaction center *pufML* and *puhA*, C4-dicarboxylate transport *dctBD*, adhesin transport system *lapBCE*, Type III secretion system, pyruvate metabolism *porABC*, multicomponent Na^+^:H^+^ antiporter *mnhBD*, glucose/mannose transport system *gtsBC*, formate dehydrogenase *fdoHI*, mannopine transport system *attA1A2BC*, Type VI secretion system, tungstate transport system *tupABC*, UDP-glucose/iron transport system *fetAB*, malonate decarboxylase *mdcABCDEG*, C1 metabolism *mgsABC mxaFI* and *mxaACGKL*, succinoglycan biosynthesis *exoALOUWY*, two-component system *adeRS*.

**Figure S6.**
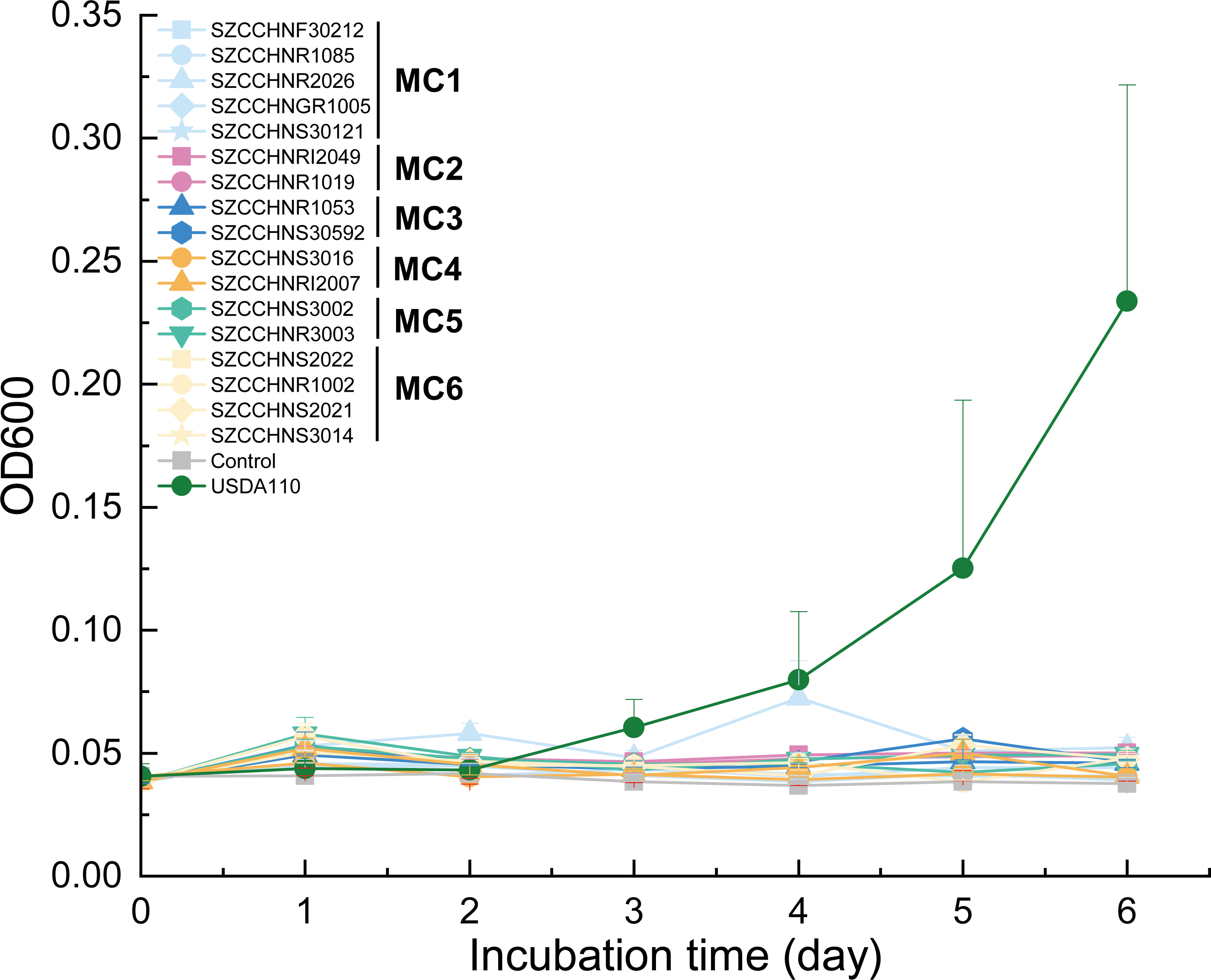
Growth of representative strains from the populations delineated by PopCOGenT for the PB of *Bradyrhizobium* on methanol as a sole carbon source, with the presence of lanthanide (Ln) species (Ce^2+^, 30 uM). The reference strain *Bradyrhizobium diazoefficiens* USDA110 was used as a positive control. Error bars indicate the standard deviation of the mean from three replicates.

**Figure S7.**
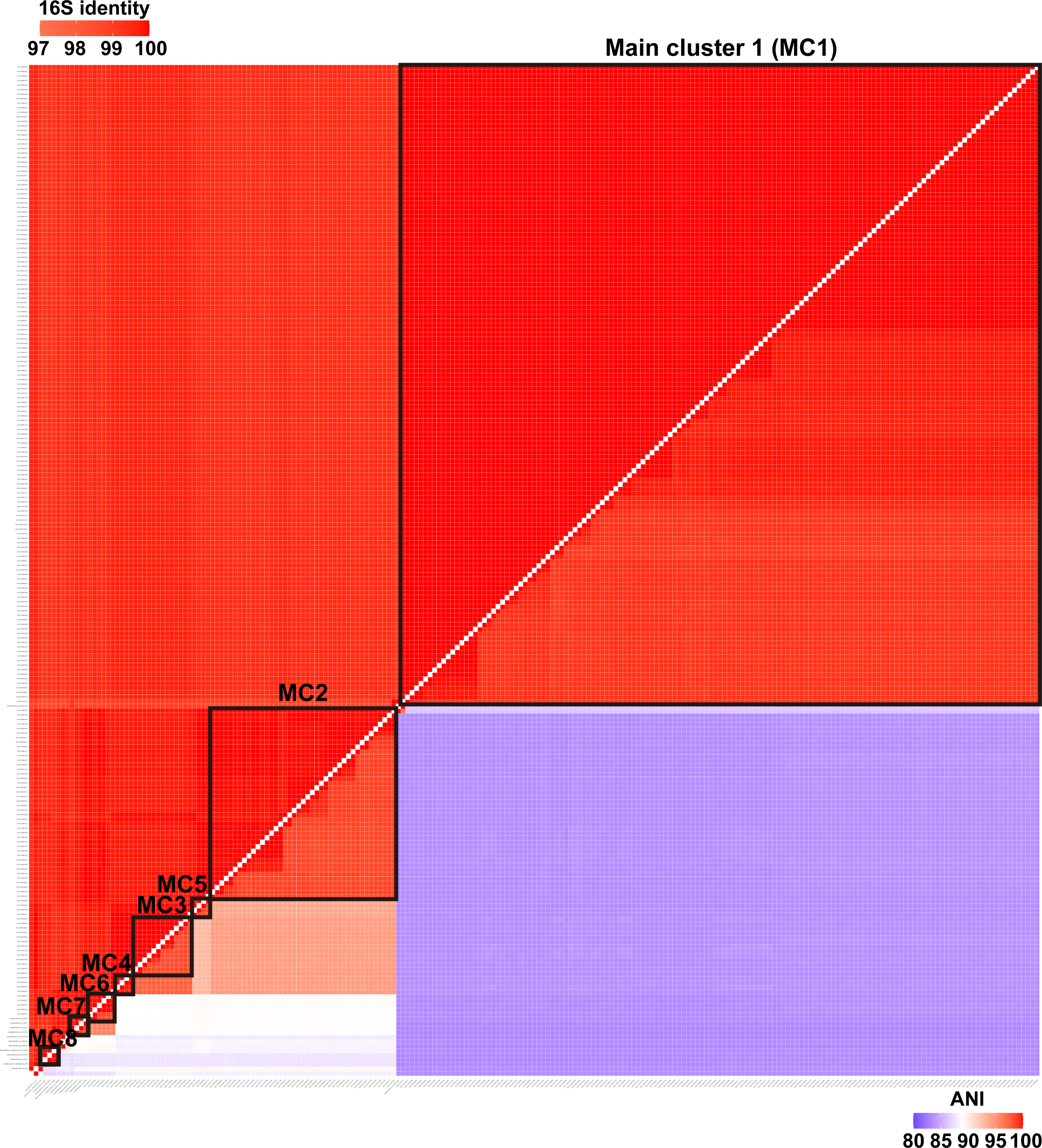
The heatmap of the pairwise identity of 16S rRNA genes and the whole-genome average nucleotide identity (ANI) of all PB members.

**Figure S8.**
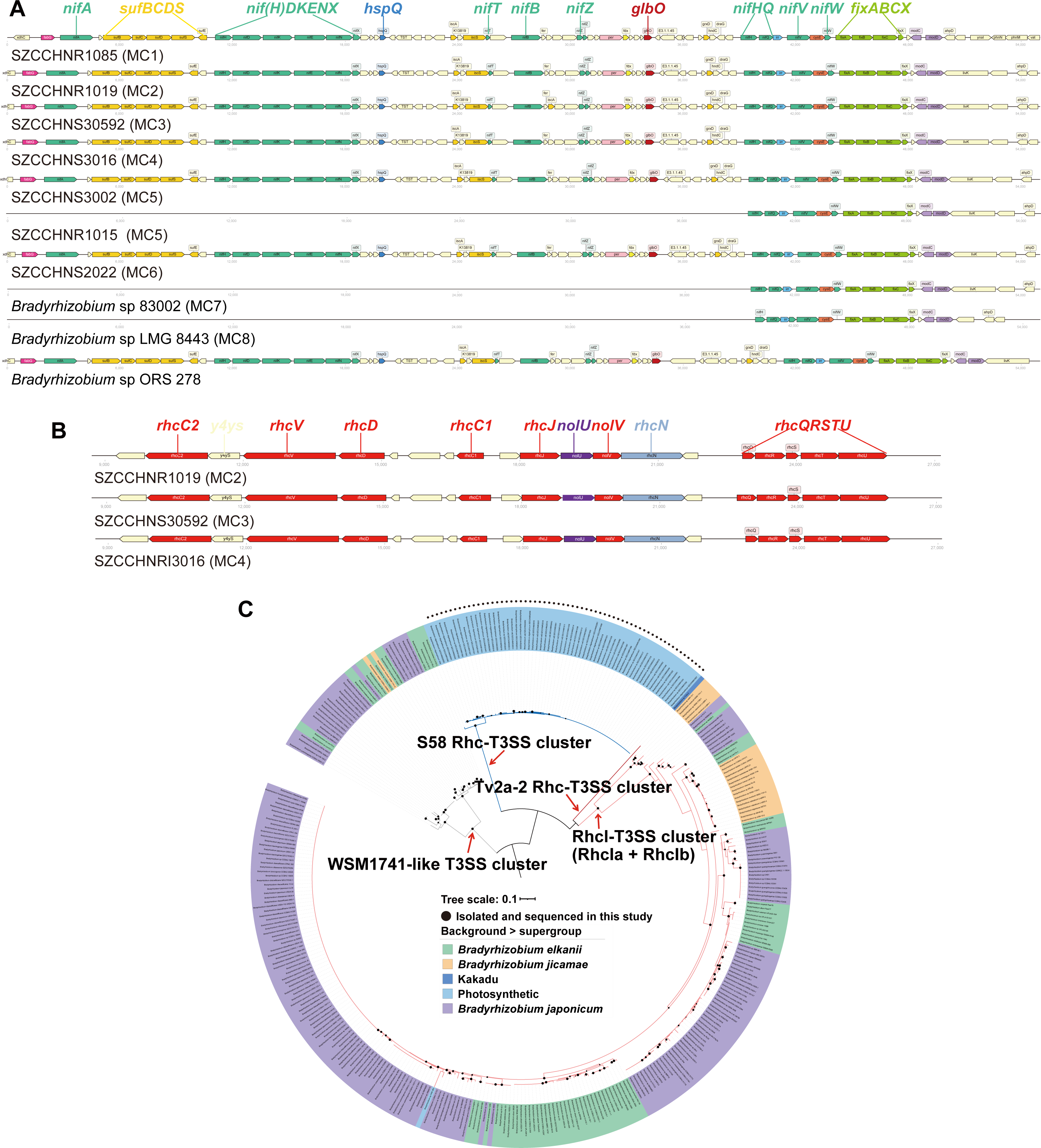
Comparison of the genomic context of the *nif* gene cluster (*nif* island) (A) and T3SS gene cluster (B) in the representative strains of PB of *Bradyrhizobium*. Gene functions are distinguished by different colors. The visualization of gene arrangement is performed with DNA-features-viewer v3.0.3 (Zulkower and Rosser, 2020). (C) The phylogenetic tree of the *rhcN* protein from *Bradyrhizobium*. The *rhcN* families were defined according to Teulet et al. (2020). The gene tree was rooted using the minimum variance (MV) method. The different colored branches correspond to the distinct genetic organization of the T3SS clusters to which the *rhcN* gene belongs. The *rhcN* in the strain *Bradyrhizobium* sp. 36 is not shown as it belongs to a different type of T3SS, likely a result of HGT from distantly related bacteria according to Teulet et al. (2020). Black circles in the phylogeny indicate nodes with IQ-Tree’s ultrafast bootstrap values ≥ 95%.

**Figure s9.**
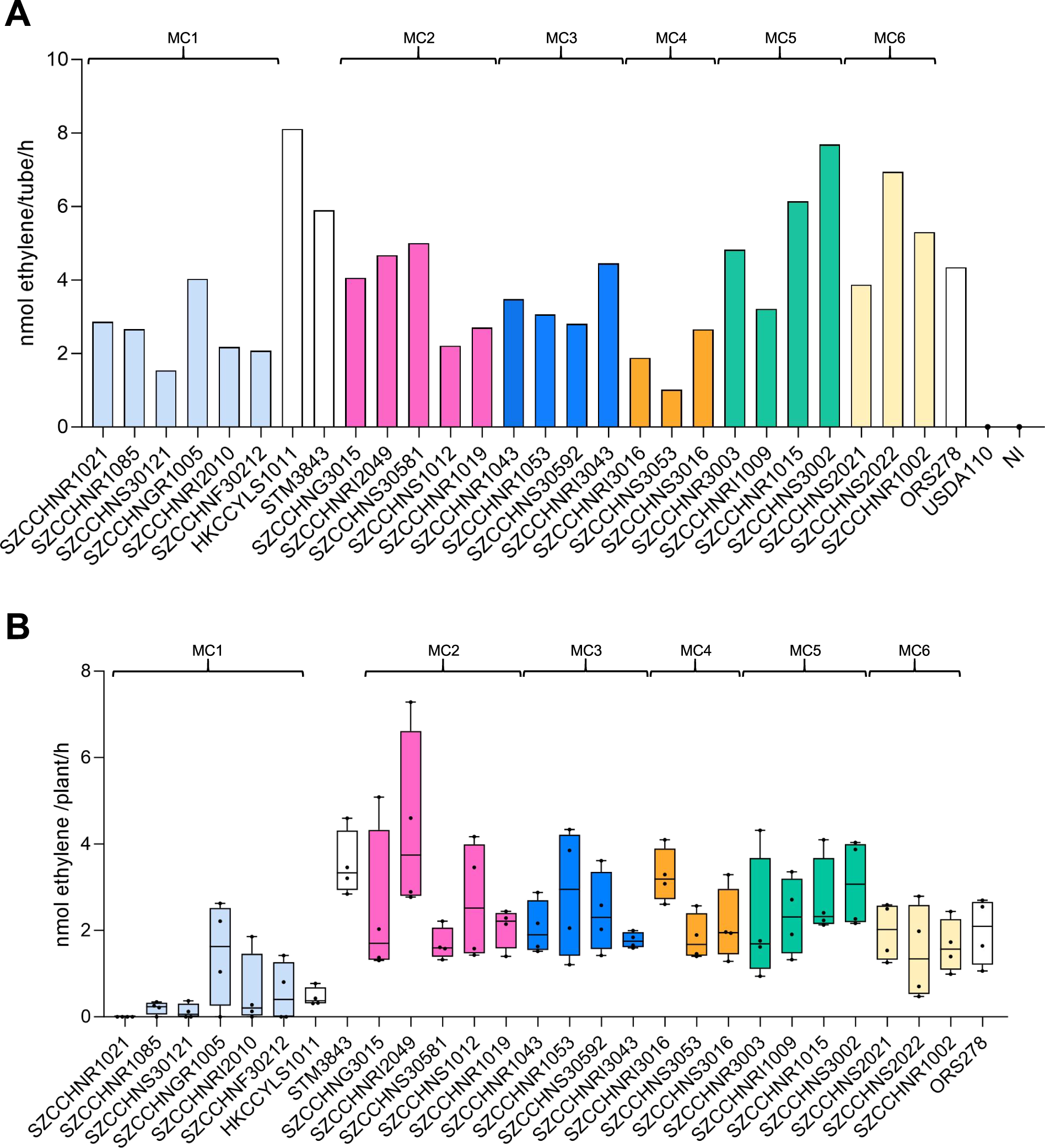
Ability of several representative strains of the main populations identified in PB supergroup to fix nitrogen during their free-living and symbiotic states. (A) Free-living nitrogen fixation after 8 days of culture in vacutainer tube. (B) Nitrogen fixation of *A. indica* plants inoculated with different PB representative strains at 17 days post-inoculation. In (A) and (B), ORS278 is used as a positive control and USDA110 (a member of *B. japonicum* supergroup) is used as a negative control in (A). NI: non inoculated. MC: Main Cluster

**Figure s10.**
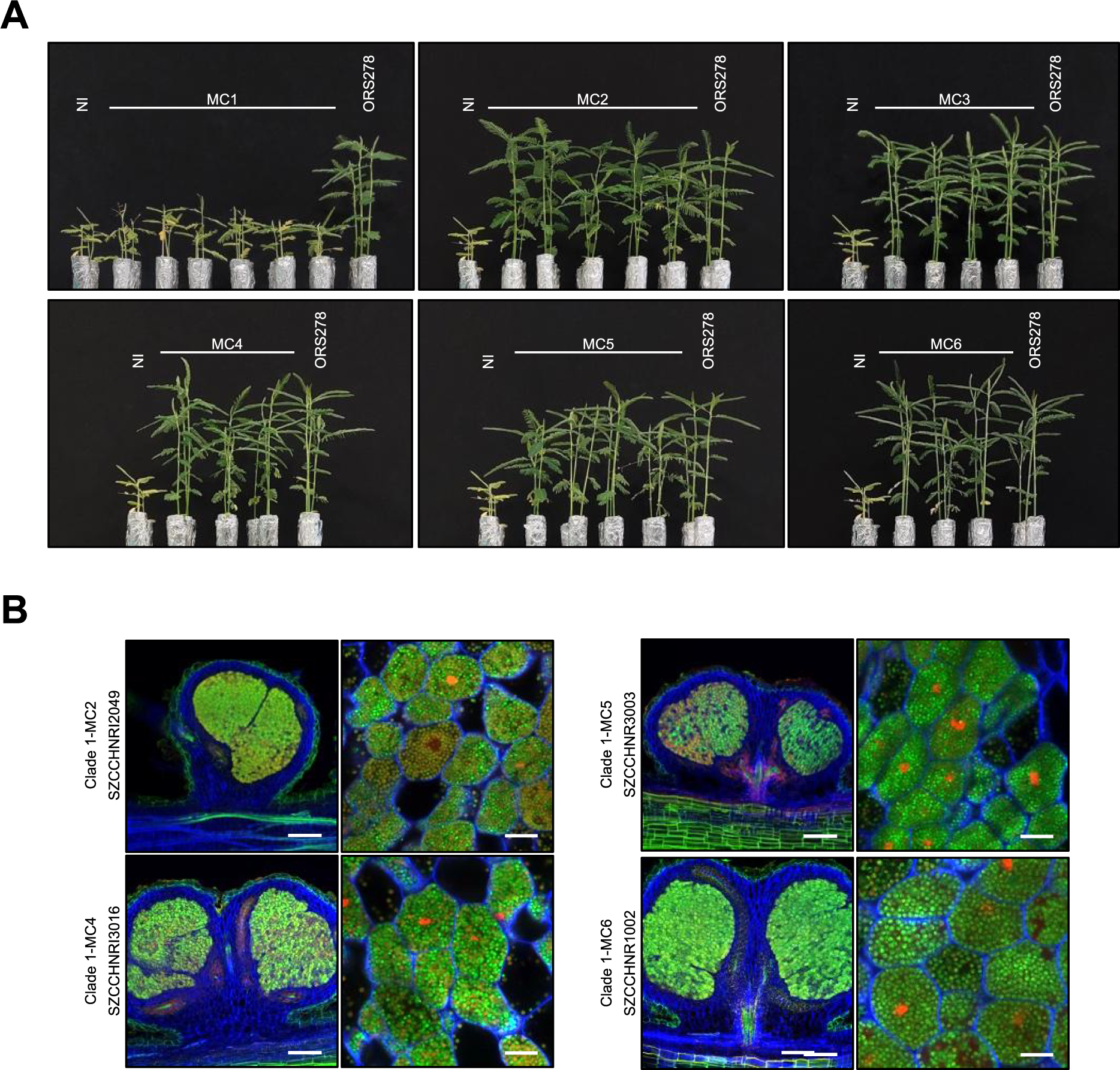
Complementary data of Fig. 2 showing symbiotic properties of several representative strains of the main populations identified in PB supergroup. (A) Comparison of the growth of the *A. india* plant (leaf phenotype) non-inoculated **(NI)** or inoculated with different representative strains of PB. All representative strains tested from each MC are present in this order: MC1 - SZCCHNR1021; SZCCHNR1085; SZCCHNS30121; SZCCHNGR1005; SZCCHNRl2010; SZCCHNF30212; MC2 - SZCCHNG3015; SZCCHNRl2049; SZCCHNS30581; SZCCHNS1012; SZCCHNR1019; MC3 - SZCCHNR1043; SZCCHNR1053; SZCCHNS30592; SZCCHNRl3043; MC4 - SZCCHNRl3016; SZCCHNS3053; SZCCHNS3016; MC5 - SZCCHNR3003; SZCCHNRl1009; SZCCHNR1015; SZCCHNS3002 and MC6 - SZCCHNS2021; SZCCHNS2022; SZCCHNR1002. ORS278 is used as control. (B) Confocal microscopy images of micro-section of nodules elicited by the other Glade 1 strains tested after staining with SYTO9 (green, live bacteria), propidium iodide (red, infected plant nuclei and dead bacteria or bacteria with compromised membranes) and calcofluor (blue, plant cell wall). Scale bars: column 1, 200 µm; column 2, 10 µm.

**Figure S11.**
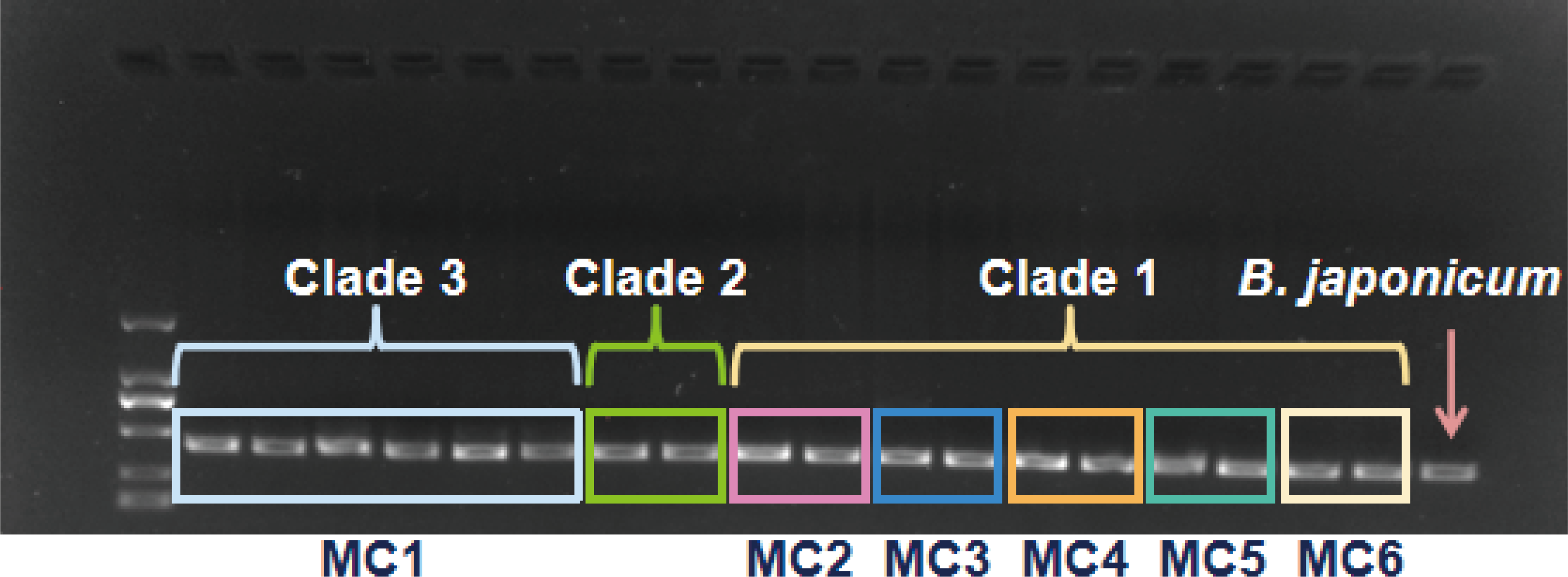
Gel electrophoresis image of DNA from PB strains amplified with the specific *rpoB* primer 2106F/BR2516R.

